# LeGenD: determining N-glycoprofiles using an explainable AI-leveraged model with lectin profiling

**DOI:** 10.1101/2024.03.27.587044

**Authors:** Haining Li, Angelo G. Peralta, Sanne Schoffelen, Anders Holmgaard Hansen, Johnny Arnsdorf, Song-Min Schinn, Jonathan Skidmore, Biswa Choudhury, Mousumi Paulchakrabarti, Bjorn G. Voldborg, Austin W.T. Chiang, Nathan E. Lewis

## Abstract

Glycosylation affects many vital functions of organisms. Therefore, its surveillance is critical from basic science to biotechnology, including biopharmaceutical development and clinical diagnostics. However, conventional glycan structure analysis faces challenges with throughput and cost. Lectins offer an alternative approach for analyzing glycans, but they only provide glycan epitopes and not full glycan structure information. To overcome these limitations, we developed LeGenD, a lectin and AI-based approach to predict *N*-glycan structures and determine their relative abundance in purified proteins based on lectin-binding patterns. We trained the LeGenD model using 309 glycoprofiles from 10 recombinant proteins, produced in 30 glycoengineered CHO cell lines. Our approach accurately reconstructed experimentally-measured *N*-glycoprofiles of bovine Fetuin B and IgG from human sera. Explanatory AI analysis with SHapley Additive exPlanations (SHAP) helped identify the critical lectins for glycoprofile predictions. Our LeGenD approach thus presents an alternative approach for *N*-glycan analysis.

**Graphical Abstract:** 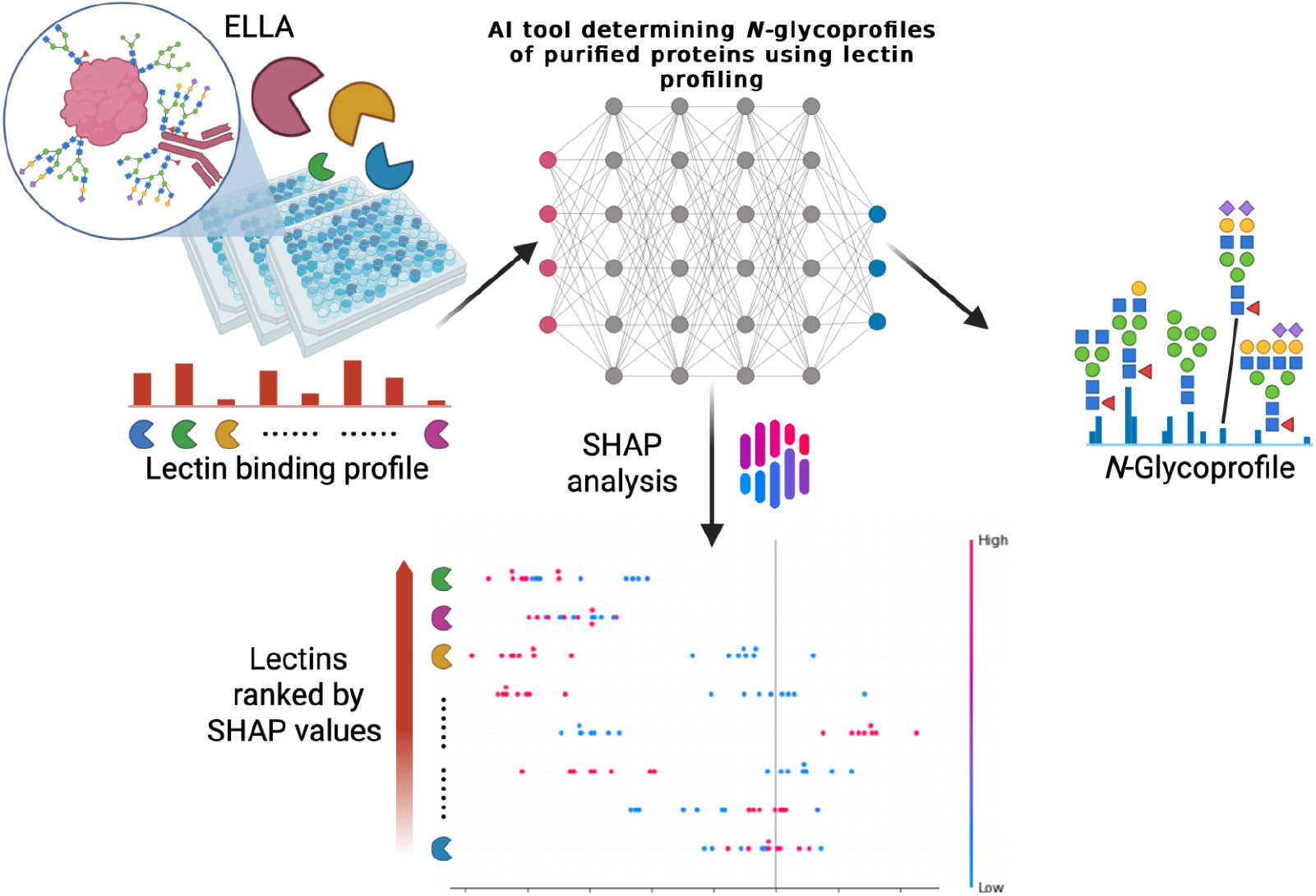

## Introduction

Glycosylation affects protein structure, function, and interactions by ensuring protein stability, proper folding, and solubility. This post-translational modification impacts diverse processes, including pathogen binding, cell adhesion, signal transduction, and molecular trafficking^1,2^. Changes in glycan structures can influence many biological and physiological processes^3,4^. For example, changes in glycosylation can modulate inflammatory responses, facilitate viral immune escape, aid metastasis in cancer, orchestrate apoptosis, and participate in the pathophysiology of various genetic and infectious diseases^5,6^. Thus, comprehensive profiling of protein glycosylation is critical to biomedical research, including the study of how single sugar residues can considerably alter protein function and activity^7–9^, and the identification of prognostic and diagnostic biomarkers for diverse diseases and genetic defects^10–12^.

Several analytical methodologies can measure glycan features, such as high-performance liquid chromatography (HPLC)^13^, capillary electrophoresis^14,15^, mass spectrometry (MS)^16,17^, nuclear magnetic resonance^18^, lectin arrays^19,20^, and enzyme-linked immunosorbent assays^21^. While each has provided invaluable information addressing specific biological questions, glycomics continues to trail behind the strides achieved by other omics, especially in next-generation sequencing (NGS) of nucleic acids^22–26^. Considering the growing awareness of the needs of glycobiology research, there is a need for newer techniques for assaying and quantifying glycan structures.

Lectins are carbohydrate-binding proteins that recognize mono- and oligosaccharide structures and can be applied to a wide variety of analytical formats^27–29^ to measure glycan features^19,30^. Despite their value for quantifying glycosylation patterns across diverse samples and conditions, lectin arrays and assays remain under-utilized, and have yet to be used to measure entire glycan structures^19^. Moreover, samples with considerably different glycan compositions can produce similar lectin profiles, making their differences difficult to observe^31,32^. Lastly, although successful work has been done in elucidating lectin binding motifs, many are not inferred while accounting for all possible glycan structures. In extreme cases, they may be restricted to single glycan structures or assays to test binding, including limited sampling of all possible glycans, or from a single type of array, and do not account for external factors^33–35^. Therefore, it is challenging to leverage lectin-based approaches for the comprehensive elucidation of intact glycoform structures in a given glycoprotein, since they do not account for many intracellular constraints that help shape the orientation of the sugars in complex glycan structures. Recognizing these challenges, there’s a clear need for more data-driven innovations.

In recent years, the incorporation of artificial intelligence (AI) models to tackle intricate biological challenges has grown prevalent. AI can help uncover hidden connections among diverse data fragments, thereby facilitating the extraction of crucial biological insights^36^. It has been demonstrated that AI models can successfully analyze complex omics data for diverse applications, such as the reconstruction of metabolic pathways^37^, drug discovery, and biomarkers^38,39^. Glycomics undoubtedly involves the deciphering process of hidden connections, especially given the ambiguity found in glycan structures (such as the non-template-directed, flexible, and repeating structural units available for a variety of modifications) and the various molecules that bind to glycans. In fact, several studies have successfully applied AI tools^40–42^ to their glycomics research. Essentially, these studies successfully extrapolated glycan-binding properties without performing costly and labor-intensive experiments by harnessing neural network (NN) models. Additionally, AI-based approaches^43,44^ are also adopted to illustrate the structure and functional relationships of glycans. Recent advances in explainable AI have now made it possible to interpret black-box AI models. SHAP (Shapley Additive Explanations),^45–47^ for instance, is a game-theoretic approach that dissects individual predictions of multifaceted AI models, allowing the computation of Shapley values for the input of essential features for local and global model interpretation. Interpretability is particularly crucial to inform and guide future decision making. The development of AI-based methods in the glycobiology field is undoubtedly burgeoning, with numerous prospects to fully unlock its capability to offer insights into the mechanisms of glycan synthesis and the interpretation of glycomic data.

Here, we present a method called LeGenD (*<u>Le</u>*ctin to *<u>G</u>*lycoprofile *<u>EN</u>*hanced with *<u>D</u>*ata-driven methods) glycoprofiling (Fig. 2), which uses an artificial neural network (ANN) model to predict *N*-glycan structures using enzyme-linked lectin assays (ELLA). Specifically, we obtained data from diverse large-scale ultra performance liquid chromatography (UPLC) glycan profiling experiments using a genetically engineered CHO cell panel. By harnessing established lectin specificities^48^, we simulated the lectin binding profiles of the 309 collected glycoprofiles. With this data, we trained the LeGenD model and validated it using measured human serum IgG and bovine Fetuin B lectin profiles. The model’s performance was evaluated by comparing their ANN-generated glycoprofiles with those derived from UPLC and high overall accuracy was achieved. Lastly, we used SHAP to shed light on the dominant lectins responsible for reconstructing the glycoprofile. Taken together, LeGenD offers an alternative platform to easily decipher comprehensive glycan structures with high-throughput and cost-effective lectin profiles. Thus, LeGenD shows increased potential of lectins for glycan analysis.

**Figure 1.**
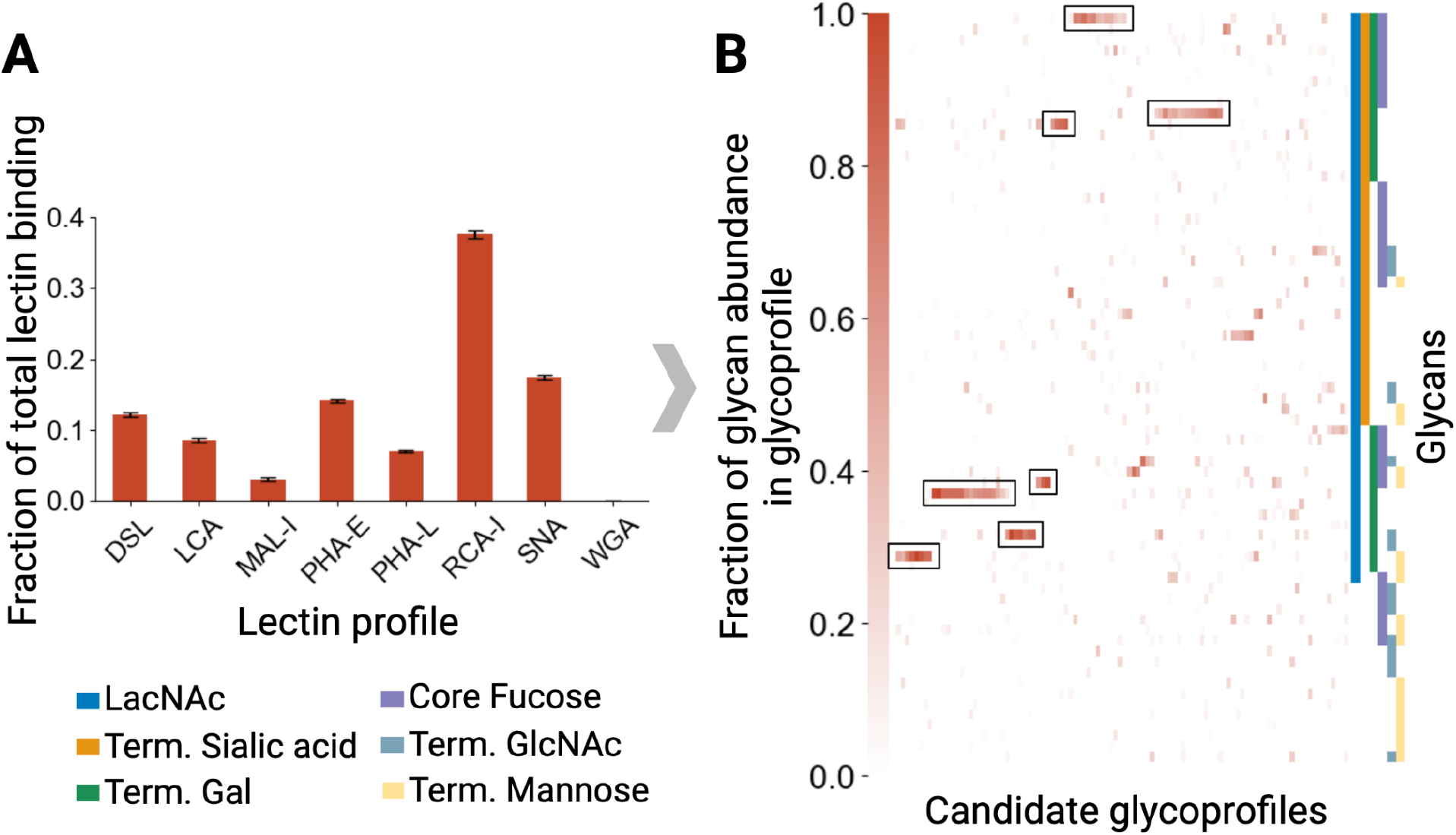
Lectin profiles cannot be connected to a single glycoprofile. From 2912 randomly generated glycoprofiles, we simulated their lectin profiles and then plotted the glycoprofiles corresponding to the 100 most similar simulated lectin profiles (Pearson R correlation > 0.95). A) Bar plot of mean lectin profiles representing their fraction of total lectin binding, with standard error bars showing the variability for each lectin. B) The heatmap shows the candidate glycoprofiles after clustering using the Voorhees algorithm. Glycan features are annotated on the right.

**Figure 2.**
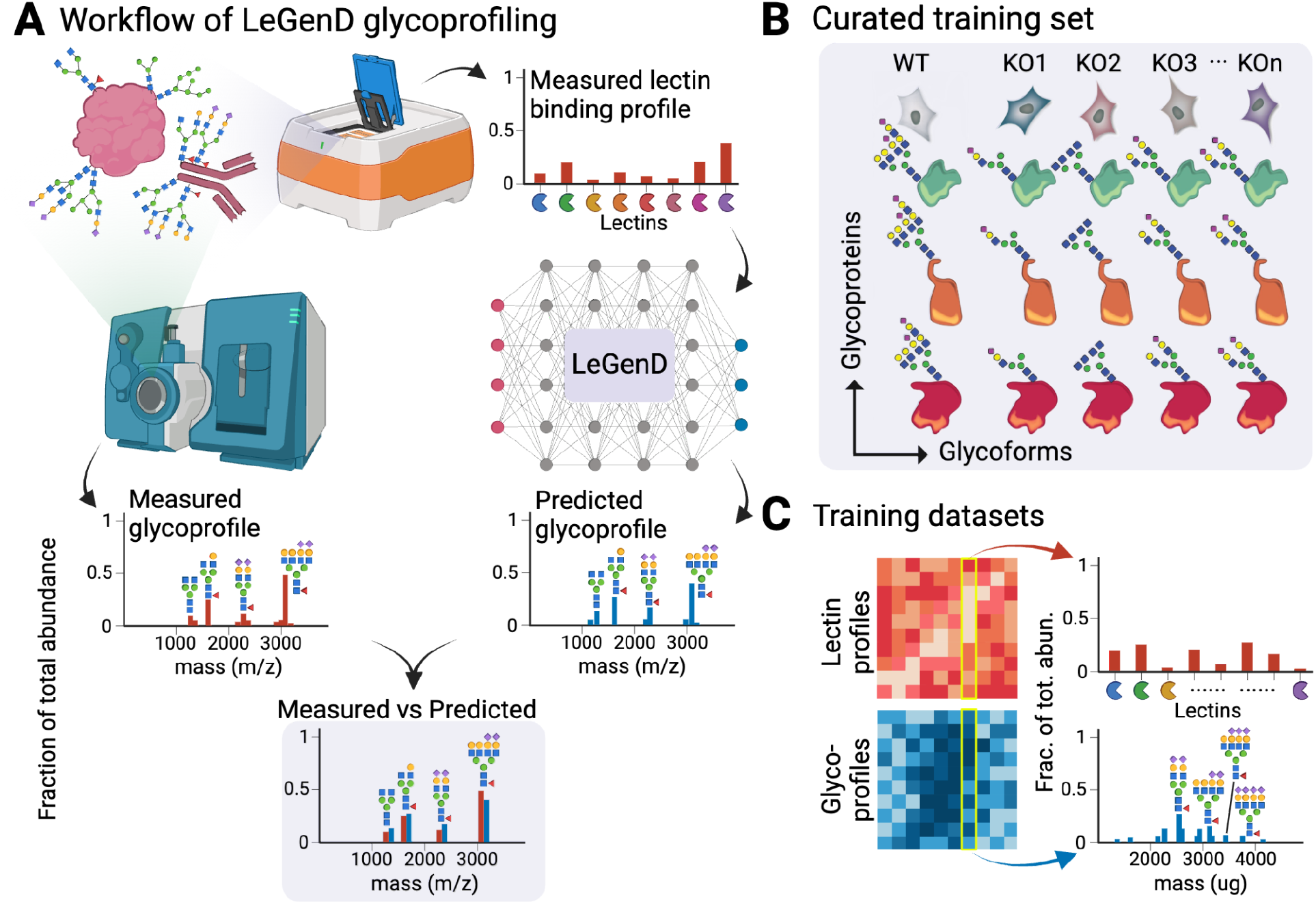
Predicting relative N-glycan abundance using machine learning. A) Workflow of LeGenD utilizing neural networks to predict N-glycoprofiles based on lectin binding profiles. B) 309 N-glycoprofiles were obtained from 10 glycoproteins expressed in a large panel of 30 glycoengineered Chinese hamster ovary cells (geCHO), yielding a comprehensive view of N-glycans for training LeGenD. C) Matrix representations of the training datasets consisting of lectin profiles and glycoprofiles.

## Results

### Lectin profiling cannot unambiguously predict glycoprofiles

Lectins are valuable reagents for measuring glycan features, but the diversity of how glycan features can be oriented in glycan structures^29,49,50^ has limited the ability to use lectins for unambiguously determining glycan structures. Furthermore, many lectins can have secondary specificities, thereby increasing the potential glycan structures they bind^51^. To estimate the scale of this problem, we aimed to quantify the diversity of glycoprofiles that could be obtained from a single lectin profile. For this, we generated 2912 random glycoprofiles from a set of 68 known glycan structures that were measured from a panel of 10 glycoproteins expressed in 30 glycoengineered Chinese hamster ovary (geCHO) cell lines (see Methods). From these glycoprofiles, we simulated their lectin profiles by multiplying a glycan-feature matrix (where each glycan is decomposed into counts of expressed features with each glycan feature annotated as per Linearcode^52^) by a binding-rules matrix (Fig 3A). This process effectively incorporated affinity factors between each glycan feature and lectin, which were derived by leveraging published^48^ glycan-lectin specificities.

**Figure 3.**
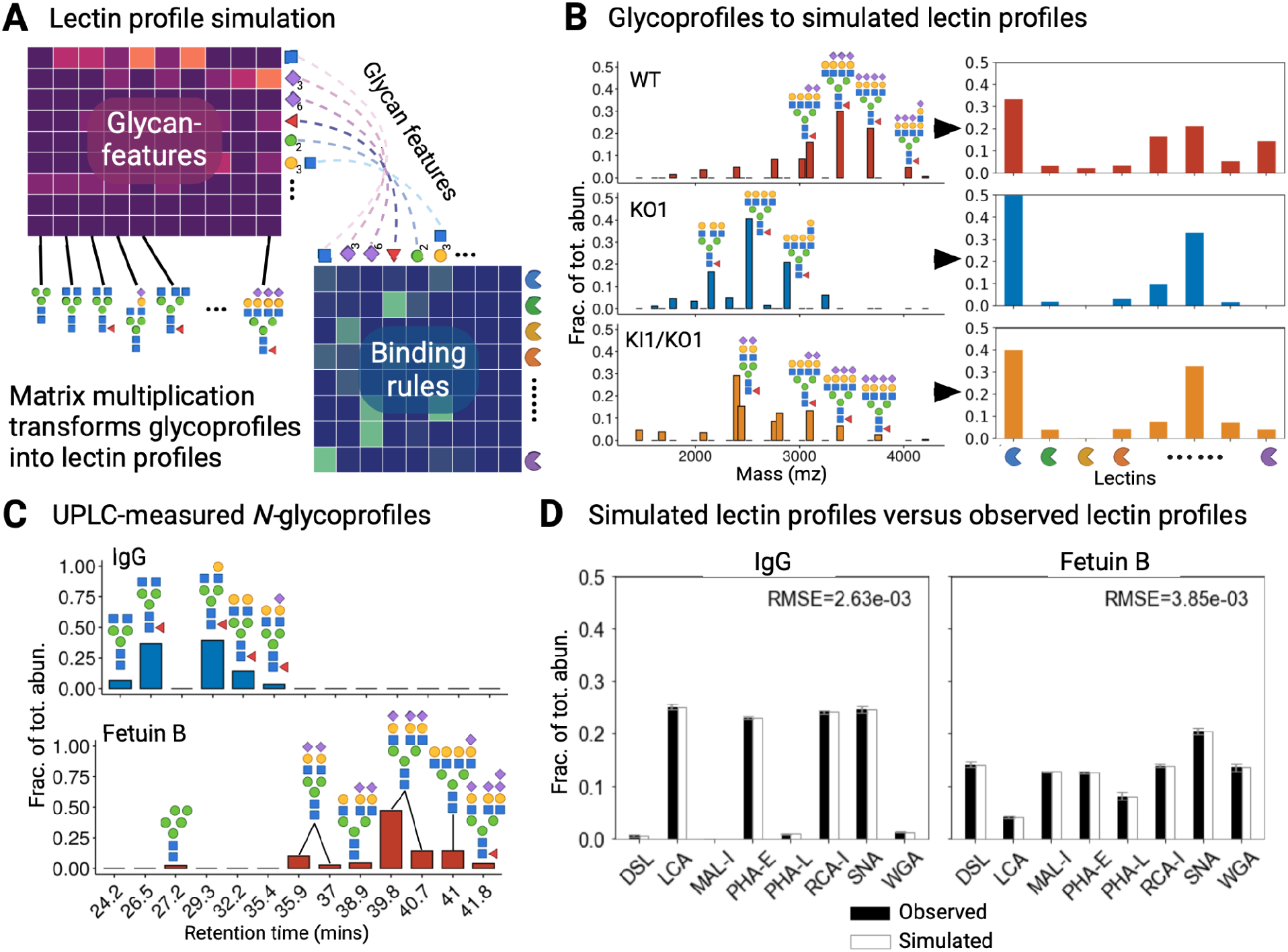
Simulating lectin profiles for model training data. A) Schematic approach for simulating lectin profiles from glycoprofiles through matrix multiplication using lectin binding rules. Glycans are mapped to lectins by decomposing each glycan into its constituent glycan features recognized by each lectin. B) Simulation of glycoprofiles to their respective lectin profiles. C) UPLC-derived N-glycoprofiles for IgG from human sera and bovine Fetuin B, Validation of simulated lectin-profiling by comparing simulated lectin profiles of IgG and Fetuin B, generated from their UPLC-derived N-glycoprofiles, D) Validation of lectin ELLA data for Bovine Fetuin B samples and IgG for human sera, generated from UPLC analysis, against actual ELLA experiments with standard error bars. The comparison reveals low RMSE values, confirming the accuracy of our simulation.

We discovered that from the 2912 randomly generated glycoprofiles, we found 100 highly-similar simulated lectin profiles (R > 0.95), clearly demonstrating that it is nearly impossible to unambiguously reconstruct glycoprofiles from lectin-based glycan features. For further insights into the diversity of glycoprofiles that can be obtained from a single lectin profile, we performed hierarchical clustering using the Voorhees algorithm, which revealed at least seven groups of potential glycoprofiles (depicted as black squares) highly correlated with the simulated mean lectin profiles (Fig. 1). However, each cell constrains the glycoprofiles they produce, e.g., based on the species, their enzyme repertoires, monosaccharide availability and structure of the receiving glycoprotein; thus, the range of possible glycoprofiles would be greatly reduced. Since the quantitative influence of many of these constraints are not well understood, we hypothesized that this can be feasibly overcome by training an ANN on highly diverse glycoprofiles, which accounts for the inherent constraints on glycan patterns imposed by its training data.

### Simulated and experimental lectin-binding profiles are consistent

LeGenD uses an ANN to connect patterns in lectin-binding to glycoprofiles (Fig. 2A). However, there is a paucity of available datasets that contain detailed lectin-binding patterns and their corresponding glycan structural information. To address this challenge, we generated simulated lectin profiles from a large study of 309 *N*-glycoprofiles from 30 geCHO cell lines, producing 10 different glycoproteins (Fig. 2B).

The simulation of the lectin binding profiles was performed in similar fashion to the preceding section (Fig. 3A). Thereafter, we aligned the simulated lectin profiles with their experimentally observed counterparts, thereby enhancing the credibility of our simulated lectin profiles (Fig. 3D).

While our simulated lectin-binding patterns are based on published binding factors, it is imperative to test if the values are consistent with our experimental data. This is especially crucial in relation to validate our assumption, that the binding intensity is proportional to the amount of glycan feature that the lectin binds to. Thus, we validated this using IgG from human sera and bovine Fetuin B as representative glycoproteins, which both contain different *N*-glycan structures^53–55^. The *N*-glycans of both glycoproteins were characterized separately using Ultra-Performance Liquid Chromatography (UPLC)^56,57^, and the relative amounts of respective *N*-glycans were estimated from the peak regions of the fluorescence chromatograms (FLC). Adhering to our simulation rules, we constructed the simulated lectin profiles for each glycoprotein. Concurrently, we employed ELLA to measure 10 distinct *N*-linked glycan features (Table S1), resulting in experimentally derived lectin profiles. Linear regression analysis indicated a strong correspondence between the simulated and experimental lectin profiles. Moreover, the analysis also provided insights into the relationship between the experimental and simulated lectin profiles, allowing us to align the simulation data with the observed data for increased accuracy. Following the application of a linear regression model, our results demonstrate a significant resemblance between the simulated lectin profiles derived from FLC data and the actual lectin profiles for both IgG (RMSE=2.63e-03) and Fetuin B (RMSE=3.85e-03) (Fig. 3B). Hence, simulated lectin profiles are highly suitable training data for LeGenD in subsequent experiments.

### LeGenD accurately predicts *N*-glycosylation in human IgG

To test the ability of LeGenD to accurately predict various *N*-glycoforms, we used 30 geCHO cell lines comprising different knockout/in combinations of 17 glycosyltransferase gene combinations to produce a range of different glycoforms of ten recombinant glycoproteins, including monoclonal antibodies, fusion proteins, cytokines, and enzymes (manuscript in preparation). From these glycoproteins, we performed *N*-glycan analysis using HILIC-UPLC profiling, which resulted in 309 glycoprofiles. We then performed lectin simulations, as described in the previous section. Thus, we obtained the training dataset and used it to configure the LeGenD architecture.

We then built a fully connected neural-network model (Fig. 4A), connecting eight lectin measurements with 68 distinct *N*-glycan structures stemming from the 309 geCHO glycoprofiles. To search for the optimal performance of the model, we performed hyperparameter tuning (see Methods) and confirmed a model architecture consisting of four hidden layers and 20 nodes in each layer (Fig. 4A) performed the best (RMSE of -3.60E-02) (Table S2). This architectural choice strikes a balance between accuracy and computational cost, as an inadequate node count in a hidden layer may limit data representation, while an excessive count might lead to overfitting during training and compromise the predictive capacity for new data^58^. The detailed hyperparameter sets that were tested are provided in Supplementary Table 2.

**Figure 4.**
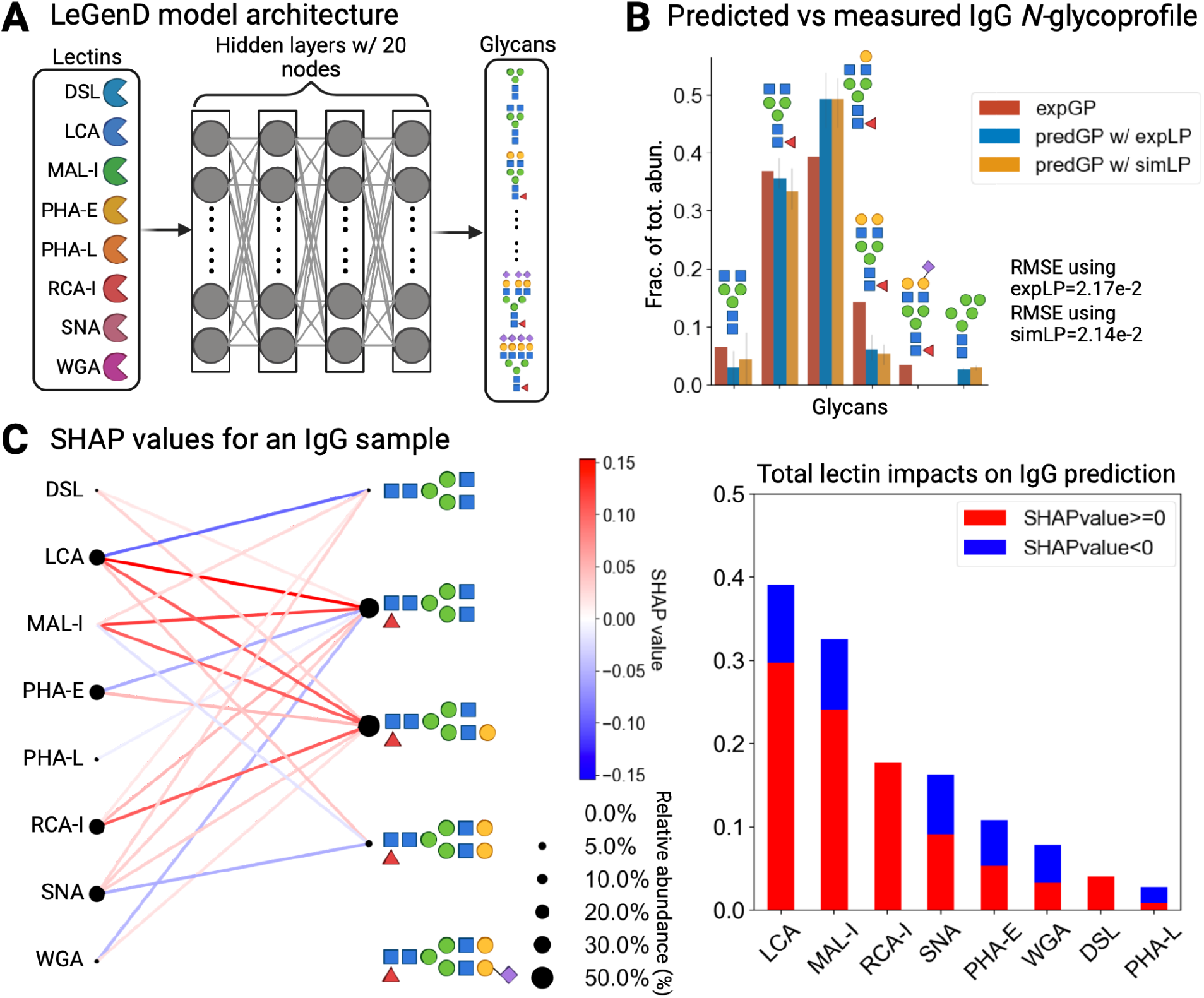
Predicting IgG N-glycoprofile with lectin binding profiles with SHAP interpretations. **A)** Architecture of the neural network model LeGenD, consisting of four fully connected hidden layers with 20 nodes in each. **B)** Comparison between the experimentally observed IgG N-glycoprofile and the LeGenD-predicted N-glycoprofile. The red bar represents glycoprofile obtained using UPLC, while the orange and blues are predicted glycoprofiles obtained by LeGenD using simulated and observed lectin assays as inputs. The fraction of total abundance for each glycan structure was also determined from UPLC-generated and lectin-based (simulated and experimental) glycoprofiles in human IgG. The error bar indicates standard error. The symbol of each glycan follows the SNFG^44^. **C)** In the bipartite network on the left, SHAP values of lectins and their glycan structures in human IgG are displayed. The color scale represents SHAP values, while the dot size indicates the relative abundance of a lectin or glycan. To the right, a bar graph breaks down the sum of the SHAP values, illustrating the contribution of each of the eight lectins used. Purple diamonds oriented towards the right and left represent ɑ-2,3- and ɑ-2,6-linked sialic acids respectively.

This model was evaluated for its ability to computationally predict *N*-glycans from ELLA-derived lectin profiles. We first provided the IgG lectin profiles. Using ANN, we predicted the identity and abundance of IgG *N-*glycans (Fig. 4B). To denoise, we employed a threshold score of 0.02, and then normalized the predictions to have a summed abundance of 1. Thus, we attained high accuracy for glycans that exhibited a fractional total abundance of 0.1 or more, with a mean RMSE of 2.17e-2. Standard error analysis yielded average values of 1.14e-3 and 1.49e-3 for predictions using measured and simulated lectin profiles, respectively, demonstrating the robustness of our approach and its ability to accurately predict *N*-glycans in unseen test samples, i.e., human IgG.

### Dirichlet Distribution-Simulated Glycoprofiles Enhanced Range of *N*-Glycans that can be Predicted

Although our method can successfully predict *N-*glycans on human IgG, our existing training dataset of 68 distinct structures can be insufficient to cover all *N*-glycans available across mammalian proteins. While adding more diverse training samples could mitigate this issue, an increased training size will require increased experimental work and higher computational cost. To expand the scope of our training dataset in this proof-of-concept without requiring additional experiments, we generated 30 synthetic *N-*glycoprofiles with a Dirichlet distribution (see Methods). The resulting glycan abundances within the simulated profiles (Fig. 5A) exhibited the expected dispersion pattern, typically manifesting 2-3 prominent peaks comparable to those observed in our experimentally observed *N-*glycoprofiles.

**Figure. 5.**
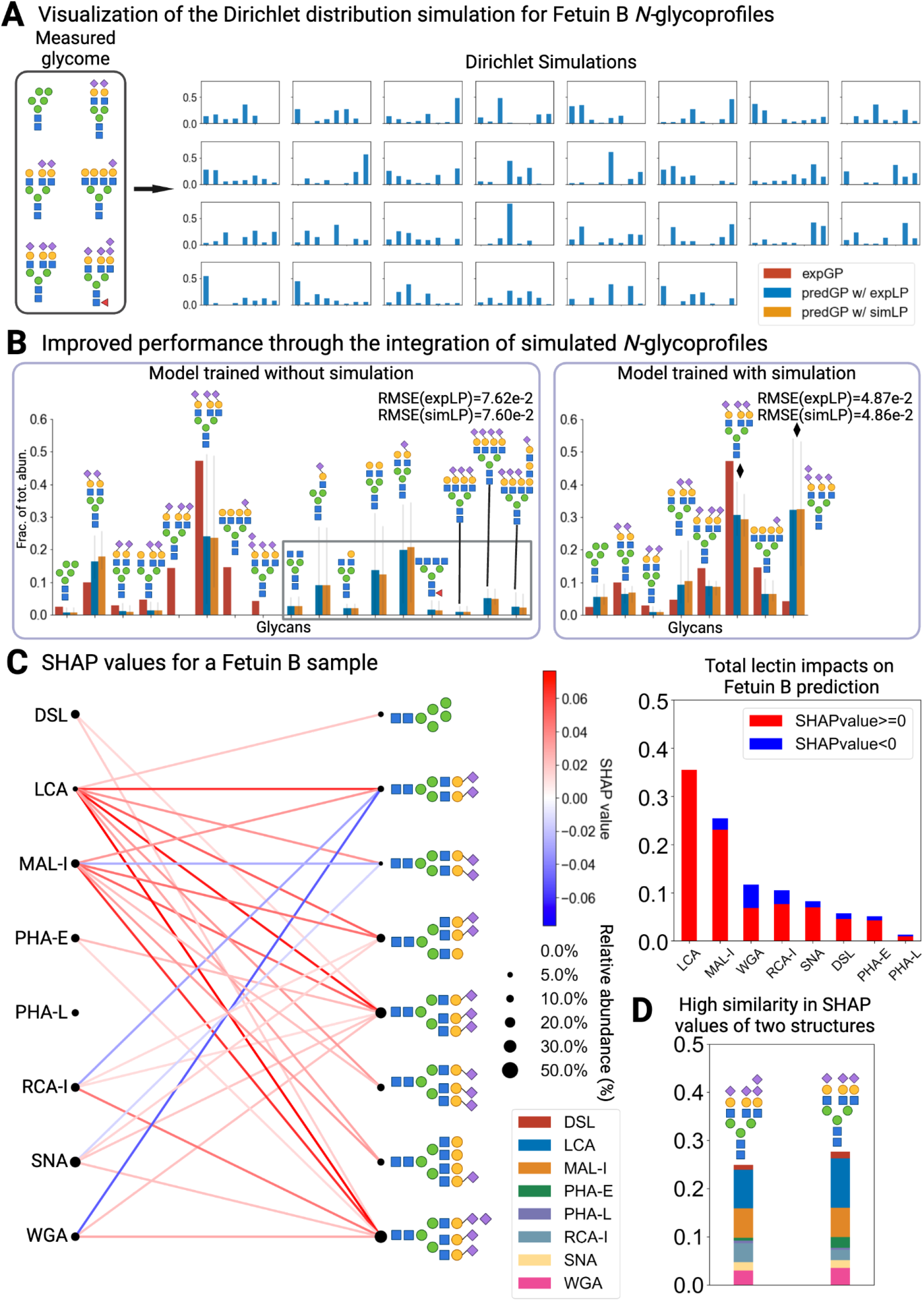
Model performance improved on unseen glycans by expand training data with simulations. **A)** To introduce unseen glycans to the model in the absence of experimental data, glycoprofile simulations were generated using the Dirichlet distribution, and then incorporated into the Fetuin B N-glycome training dataset **B)** Predictions made by LeGenD before and after the simulated Fetuin B N-glycoprofiles were added into the training dataset. Prediction accuracy got improved after this addition. **C)** Bipartite network shows the SHAP value of the lectins on the glycans expressed in the Fetuin B sample. The color scale represents SHAP values, the dot size indicates relative abundance or a lectin or glycan. **D)** Complex N-glycan structures exhibited remarkable similarity in SHAP values despite varying sialylation patterns. Purple diamonds oriented towards the right and left represent ɑ-2,3- and ɑ-2,6-linked sialic acids respectively.

Subsequently, we incorporated the 30 DD generated glycoprofiles into our model. Because this constituted approximately 10% of the original training dataset, it provided new candidate *N-*glycans while ensuring that the training process remained unbiased. We examined the accuracy between datasets consisting solely of experimentally observed glycoprofiles (ex-GPs) and combined simulated/experimental glycoprofiles (comb-GPs). Training was iterated ten times after loading the datasets into the model, and the resulting average predictions were plotted using a 0.02 threshold. We found that the model-predicted glycoprofiles using experimental lectin profiles (predGP w/expLP) and trained with the comb-GP datasets exhibited notably reduced Root Mean Square Error (RMSE) values in comparison to the ex-GP dataset (ex-GP:7.62e-2, comb-GP:4.87e-2), resulting in a reduction of false positive instances and noise (Fig. 5B: black and gray rectangles). Notably, the model-predicted glycoprofiles using the simulated lectin profile (predGP w/simLP) and trained with the comb-GP datasets also exhibited RMSE values comparable to those of the ex-GP dataset (ex-GP:7.60e-2, comb-GP:4.86e-2). We also identified inconsistencies between the quantitative relative abundances in the experimental glycoprofiles and the predicted values for tri- and tetra-sialylated *N-*glycans (Fig. 5B). This is likely due to the sialic acid-targeting lectins used: SNA and MAL-I (SNA targets ɑ-2,6-sialylated LacNAc^48^ and MAL-I targets ɑ-2,3-sialylated LacNAc^48,59^). However, neither lectin exhibited specificity for ɑ-2,8 polysialylated LacNAc epitopes^60^. Thus, the model cannot differentiate between the triantennary *N-*glycans with three terminal sialic acids and the *N-*glycan with polysialic acid based on the lectins provided in Fetuin B (Figure 4B, black diamonds). This is supported by the high similarity in the SHAP values of lectins between these two glycans (Fig. 5C bottom right) using SHAP analysis. The similar predicted abundances of both glycan structures implies that our model weighted both glycan moieties equivalently. Although these findings demonstrate that the model’s learned embeddings adapted to the simulated training data, it should also be emphasized that the chosen lectins impact the model’s ability to accurately distinguish the nuances of glycan structures. Overall, by integrating the Dirichlet distribution-based simulation of *N*-glycan glycoprofiles, we demonstrated the resilience of the model in handling previously unencountered data instances, thereby reducing the necessity for further training and parameter fine-tuning.

### Crucial Lectins can be identified using SHAP

To interpret the “black box” model in LeGenD, we employed SHapley Additive explanations (SHAP) analysis^45–47^ (see Methods). This elucidated specific lectins and their associated glycan epitopes, which contributed to the prediction of glycoprofiles using lectin profiles acquired through ELLA. The Shapley importance values for a given feature can be understood as the average incremental contribution that the feature offers to the model considering all potential groupings, in our case, the set of lectins chosen. For IgG (Fig. 4C), SHAP revealed that LCA and MAL-I were the most important lectins, with SHAP importance values of +0.3 and +0.25 above the average prediction, respectively. In contrast, RCA, SNA, and PHA-E, while still important, contributed less than +0.2 above the average prediction. In Fetuin B (Fig. 5C), a similar ranking was observed for LCA and MAL-I based on the aggregate sum of SHAP values. In addition, SHAP analysis underscored the importance of SNA and PHA-E in predicting glycan structures in Fetuin B.

While LCA and MAL-I are good predictors for the glycan structures in both proteins, both lectins exhibited differences in their interpretability, based on the different sign of SHAP values of features for the lectin-binding profiles presented to the model. This is particularly evident when examining the contribution of LCA towards the predicted glycoprofile of IgG. LCA binds to core-fucosylated *N*-glycans^48,61,62^, commonly found on IgG^63–66^. Thus, the positive SHAP values for LCA, in conjunction with core-fucosylated *N*-glycans, suggests a positive correlation between the LCA lectin-binding profile and model-predicted core fucosylation. The contribution of MAL-I towards the LeGenD-predicted Fetuin B glycoprofile can be similarly interpreted, given its affinity for terminal Siaα2-3Galβ1-4GlcNAc^48,59,67^ found on Fetuin B^68–70^.

In contrast, elevated positive SHAP values were also seen for LCA and MAL-I when predicting the glycoprofile from the lectin binding profiles of glycoproteins lacking their preferred glycan epitope. Specifically, the LCA binding profile exhibited a positive correlation (as indicated by the positive SHAP values) with the model-predicted glycoprofile of Fetuin B, despite Fetuin B not possessing core-fucosylated *N*-glycans^70^. Similarly, in the absence of the terminal Siaα2-3Galβ1-4GlcNAc epitope in IgG^71^, the MAL-I binding profile still demonstrates a positive correlation with the model-predicted glycoprofile. These results suggest that the strong positive correlation of LCA and MAL-I binding profiles with non-existent glycan epitopes is due to the lectin binding measurements being absent when their respective glycan epitopes were absent. This interpretation serves to recontextualize their contribution to the predicted glycoprofile as a negative control, as further supported by analyses wherein the exclusion of LCA and MAL-I yielded significantly less accurate predictions (Fig. S1).

## Discussion

In this study, we presented LeGenD, a method to map lectin-binding profiles to predict the identities and quantities of each *N-*glycan in a sample. We demonstrated that this approach is feasible after the model is trained with diverse glycoproteins, suggesting that the model captures the inherent relationships between glycans determined by the cell. We also found that explanatory AI approaches such as SHAP can be used to identify the most informative lectins. Thus, this study showed that LeGenD could be an alternative glycoprofiling approach that can be implemented in different formats to enable miniaturization and high-throughput glycoprofiling.Our model also provides a platform for further research and development to obtain more precise carbohydrate-binding proteins for quantitative measurements. This is particularly valuable to complement current analytical methods, which can encounter challenges in deciphering glycan structural features due to challenges from isobaric monosaccharides, stereochemistry^72,73^, and the need for comprehensive preliminary structural assignments and subsequent validation using orthogonal technologies implemented by experts in glycoanalytics^30,74^. Lastly, in the current landscape, there is a clear lack of platforms for rapidly tracking predominant glycosylation changes across diverse glycomic contexts. However, LeGenD is well poised to effectively address these challenges, thereby mitigating bottlenecks in glycan structural analysis.

LeGenD presents an alternative approach for glycoprofiling to complement the existing methods. Mass spectrometry provides precise mass/charge-based measurements of glycans and their fragments, and liquid chromatography helps identify glycans based on retention time. Lectins are particularly useful for identifying glycan epitopes and for linkage stereochemistry. Although several studies have attempted to profile glycans holistically^75^ or to predict single glycans from lectin binding patterns^33^, mapping these patterns to an actual glycoprofile is a highly under-constrained problem, wherein nigh infinite glycoprofiles could have the same lectin profile (Fig. 1). Therefore, we presented an approach to overcome this challenge by using AI with a large glycomics training set to successfully compute an *N-*glycan profile based on a lectin-binding profile. Thus, this method takes the best of different techniques, such as quantitative measurement of structural composition (chromatography and mass spectrometry) and linkage information (lectin arrays). We anticipate that LeGenD could be implemented towards other major glycosylation types (*O-*glycans) and in different formats, such as with lectin arrays^76–78^, microfluidics^79^, or Next Generation Sequencing^80–82^ to increase its throughput. However, each format will require their own training dataset.

Some lectins perform more poorly in LeGenD, as suggested by the SHAP analysis. For example, SNA exhibited a high signal-to-binding threshold with an average raw OD value of 1.22 in IgG, even though ɑ-2,6-linked Neu5Ac accounted for just 3.39% of the relative abundance in the overall *N*-glycome. Furthermore, SHAP demonstrated that SNA failed to contribute to the prediction of its preferred epitope, Siaα2-6Galβ1-4GlcNAc, which is typically present in IgG in the Fc region^71^. In the case of Fetuin B, SNA surprisingly displayed negative SHAP values towards glycans containing Siaα2-6Galβ1-4GlcNAc epitopes but positive values towards glycans containing Siaα2-3Galβ1-4GlcNAc. We anticipate that this is due to the previously observed variability in SNA measurements^83^, likely due to its formation of octomers and oligomerization given it is also heavily glycosylated^84^. Given that many lectins are glycosylated, we anticipate that the quantitative precision of LeGenD will improve with the inclusion of nonglycosylated lectins or other carbohydrate-binding molecules.

SHAP provided further insights into glycan epitopes that require better reagents for quantification. For example, we found that PHA-E and PHA-L lectins, which target complex epitopes such as bi-, tri-, and tetraantennary structures^48^, did not significantly improve the predictive performance of LeGenD for IgG and Fetuin B. We hypothesized that, while these lectins can recognize various structures, such as internal complex *N*-glycan structures, there is a need for more specific carbohydrate-binding proteins to enhance the model’s overall performance. In relation to this, another possible explanation for the limited impact of PHA-E and PHA-L on LeGenD’s predictive performance is linked to the findings of Yom et al.^33^, who analyzed lectin-glycan binding patterns. They determined that lectins exhibit a stronger affinity towards terminal glycan motifs because they are least susceptible to steric interference. Therefore, future studies are needed to identify and screen proteins with better binding specificities to the internal structure of glycans.

LeGenD can reveal local glycan feature relationships and thereby accurately profile common glycan structures and their relative abundance. However, to enhance its versatility, generating additional training data from multivariate probability distributions (e.g., Dirichlet distributions) is beneficial. This extends the inclusion of a broader array of glycan structures including bisecting GlcNAc, non-core fucose, modified sialic acids, polysialic acids, and *O*-glycans. Although this study was confined to the *N*-glycan structures from standard glycoproteins (human serum IgG and bovine Fetuin B) to recapitulate closely related glycan features from our training dataset, the framework can easily be expanded to encompass *O*-glycans and other glycoproteins of interest. Consequently, a more comprehensive exploration of the diverse glycan structures will further enhance their precise predictions.

In the present study, we demonstrated that ANNs can be used to identify an accurate glycoprofile from a lectin-binding profile, despite the immense diversity of glycans that could exist based solely on known enzyme functions. Glycans are highly complex, highly branched, nonlinear biopolymers that are synthesized through seemingly stochastic interactions between glycosyl hydrolases and glycosyltransferases^85–87^. However, it is anticipated that the physicochemical properties of the cell and the glycosylated proteins will limit the range of possible glycan structures and possibly which ones are more likely to co-exist in any given sample. Our study shows that ANNs capture these constraints on glycan synthesis, thus enabling the accurate determination of the correct glycoprofiles despite the expected theoretical complexity. However, while conventional neural networks have been successful here and in other studies for studying glycosylation patterns and processes ^88–91^, further developments in A.I. have presented new model topologies that could further refine technologies, such as ours, for more accessible technologies in glycobiology. For example, glycans have proven to be well-suited for graph neural network (GNN) applications^42,43,92,93^. Indeed, GNNs leverage the full molecular structure or even the geometry for inputs, and the GNN itself learns informative molecular representations to predict the given target properties^94–96^. In the context of glycans, they can comprehensively account for monosaccharides and their linkages, represented as nodes and edges, respectively, to form “glycowords”^93,97^. This strategic representation facilitates the capture of unique structural features and contextual information pertaining to glycans including branching substructures, anomericity, and various types of linkages^97^. GNNs have been successfully employed for motif exploration within glycan substructures, enabling categorization by *O*-/*N*-linkage, immunogenic properties, evolutionary origin, and viral protein recognition^43,93^. Therefore, we anticipate that integrating alternative model architectures beyond standard neural networks will further enhance applications for studying glycan structural patterns. These results are highly compatible with our workflow, resulting in an improved prediction performance.

In this proof-of-concept study, we demonstrated the ability of LeGenD to accurately estimate the glycoprofiles of *N*-glycans on two model proteins. Although our training dataset included multiple proteins with diverse properties, we only assessed *N*-glycans from proteins produced in CHO cells. Glycans from other mammalian cell types can be more complex, and other classes such as *O*-glycans are also found on thousands of proteins^6,98^. This intrinsic variability across species poses an additional challenge for comprehensive predictive modeling. The enhancement of this form of generalizability can be achieved by incorporating future samples containing an even broader spectrum of glycans and a more extensive panel of lectins.

In summary, as awareness of the importance of glycosylation increases, new technologies are needed to increase the accessibility and throughput of glycan analysis. LeGenD provides a complementary concept that with further development could aid in the proliferation of glycoanalytics in routine biological research. Indeed, leveraging AI in glycoanalytics can help manage the complexity of this field, thereby enabling a much larger user base for glycomics and allowing researchers to link changes in glycosylation with phenotypic changes.

## Methods

### *N*-glycan release and preparation from glycoproteins for LC-based glycan characterization

IgG (from human serum; Sigma-Aldrich, St Louis, MO, USA) and fetal bovine Fetuin (Sigma-Aldrich) were each treated with a mixture of PNGase F (NEB, Boston, MA, USA) following the manufacturer’s protocol.

### IgG and Fetuin B *N*-glycan purification and procainamide labeling

The *N*-glycans obtained by enzymatic treatment of fetuin and IgG were purified using C18 Sep-Pak® columns. To eliminate salts and hydrophilic species, the purified samples were subjected to an additional cleanup step using graphite cartridges. The C18 Sep-Pak column was initially conditioned with 1 ml of 10% aq MeOH + 1% HOAc three times, followed by sequential addition of 50% aq MeOH, 100% MeOH, and Chloroform. Subsequently, the column was washed twice with 1 ml of 100% MeOH, 50% aq. MeOH, and 1 ml of H2O five times. Similarly, HyperSep Hypercarb PGC columns were conditioned with 1 ml of 1% trifluoroacetic acid (TFA), 50% aq acetonitrile (ACN), and 100% ACN three times, followed by 1 ml 50% aq ACN twice, and with 1 ml H2O three times. The Sep-Pak and PGC columns were further washed with 1 and 3 ml of H2O, respectively. Finally, *N*-glycans were eluted using 3 ml of 30% aqueous ACN containing 0.1% TFA, and the elution fraction was collected and lyophilized for further analysis.

The reconstituted *N*-glycans were labeled with procainamide via reductive amination, as described by Xie et al^99^. Briefly, a procainamide solution was prepared by dissolving procainamide in sodium cyanoborohydride in a DMSO:acetic acid:H2O (280:120:100 v/v/v) solution and mixed thoroughly. Labeling solution was added to each sample and incubated at 65°C for 1 h. ACN was mixed with each sample, and SPE cleanup was performed using GlycoClean S-Cartridges (Prozyme, Hayward, CA). After cleanup, the samples were washed thrice with acetonitrile. Finally, the procainamide-labeled *N*-glycans were eluted with H_2_O.

### UPLC analysis of *N*-glycans from IgG from human sera and fetal bovine Fetuin B

*N*-glycan analysis was performed using a Waters Acquity UPLC system attached to a fluorescence detector (Besson Scientific, San Diego, CA, USA). A known amount of *N*-glycan was tagged with the fluorophore Procainamide and profiled on a (BEH Glycan, 150 mm x 2.1 mm x 1.7um). A gradient mixture of solvents containing A:100mM ammonium formate (pH-4.5) and B: acetonitrile was used to profile glycans. The solvent gradient conditions used are listed in Table 1. Detection was performed using a fluorescent detector set at excitation and emission wavelengths of 310 and 370 nm, respectively. The column temperature was set at 60°C and the autosampler temperature was maintained at 10°C.

**Table 1:**
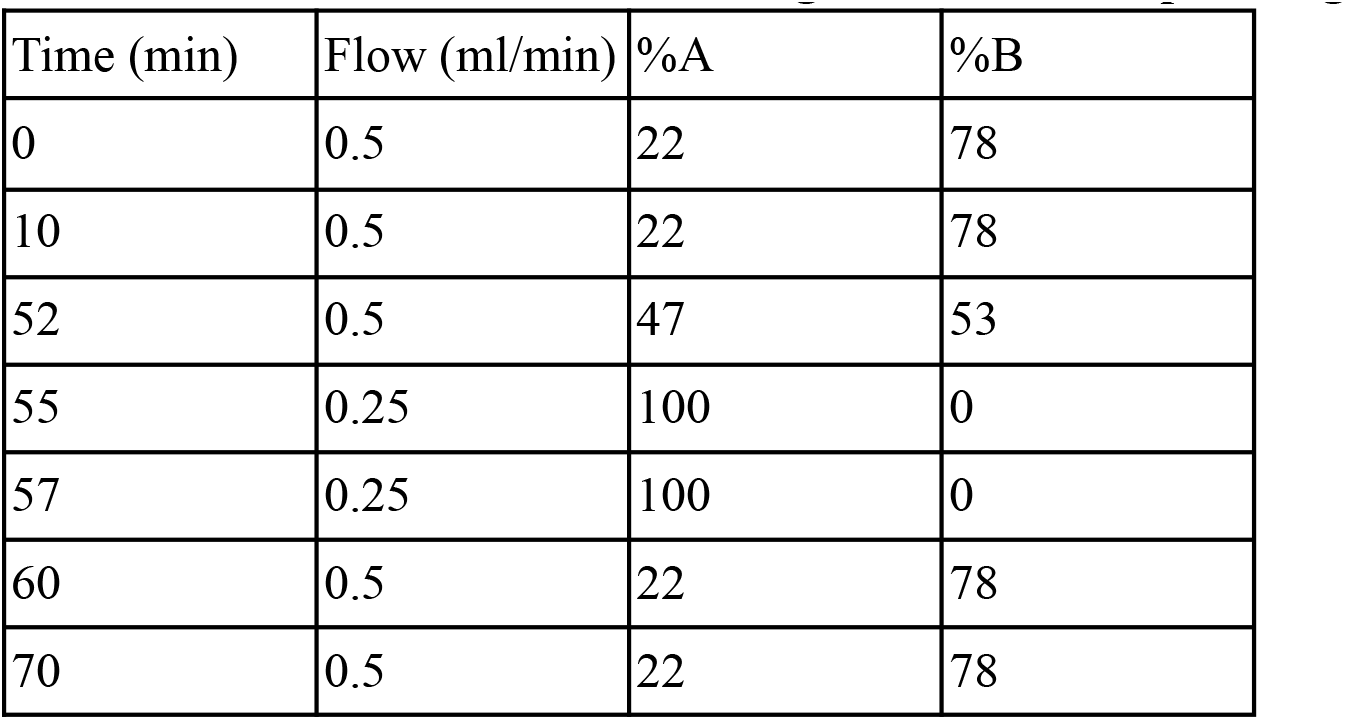
UPLC solvent mixtures for IgG and Fetuin B profiling.

### CHO-derived recombinant protein purification and HILIC-UPLC *N*-glycan analysis

Herceptin, Rituximab and Enbrel were purified by protein A affinity chromatography. For each protein glycoform, a 1-mL MAbSelect Extra column (Cytiva) was equilibrated with 5 column volumes (CV) of 20 mM sodium phosphate, 0.15 M NaCl, pH 7.2. Next, 30-mL supernatant was loaded, the column was washed with 20 CV of 20 mM sodium phosphate, 0.15 M NaCl, pH 7.2, and the protein was eluted using 0.1 M citrate, pH 3.0. The elution fractions (0.5 mL) were collected in deep-well plates containing 100 µL of 1 M Tris at pH 9 per well.

Alpha-1-antitrypsin, butyrylcholinesterase, protein Z, serpin A5, serpin A10, serpin C1, and EPO, all C-terminally tagged with the HPC4 tag (amino acids EDQVDPRLIDGK), were purified over a 1-mL column of anti-protein C affinity matrix according to the manufacturer’s protocol (Roche, cat. no. 11815024001). 1 mM CaCl2 was added to the supernatant, equilibration buffer, and wash buffer, respectively. The proteins were eluted in 0.5 mL fractions using 5 mM EDTA in the elution buffer.

For all proteins, elution fractions containing the highest protein concentration were concentrated using Amicon Ultra centrifugal filter units (MWCO 10 kDa). 12 µL of concentrated protein solutions (concentrations varying between 0.1 and 1 mg/mL) were subjected to *N*-glycan labeling using the GlycoWorks RapiFluor-MS *N*-Glycan Kit (Waters). Labeled glycans were analyzed by HILIC-FLR using an ACQUITY UPLC Glycan BEH Amide column (2.1 x 150 mm, 1.7 µm, Waters) mounted on an Ultimate 3000 UPLC system and a Fusion Orbitrap mass spectrometer from Thermo Scientific. Acetonitrile (100%) and ammonium formate (50 mM, pH 4.4) were used as mobile phases.

### Enzyme-linked lectin assay

96-well high-bind microplates (Corning, Corning, NY, USA) were coated overnight at 4°C with either 100 μl of IgG or 10 μg/mL fetal bovine Fetuin B using carbonate coating buffer (100mM NaHCO_3_, 30 mM NaCO_3_, pH 9.5). The plate was then blocked with a blocking buffer (1X PBS with 2% w/v polyvinylpyrrolidone) for 1 h at 37°C. Biotinylated lectins (Vector Labs, Burlingame, CA, USA) were diluted to 10 μg/ml in binding buffer solution (10mM HEPES, 0.15M NaCl, 0.1mM CaCl_2_, pH 7.5) and added to the plate for 30 min at room temperature. The avidin-biotinylated HRP complex (ABC reagents; Vector Lab, Burlingame, Calif., USA) was subsequently added according to the manufacturer’s instructions. Each step was washed thrice with PBST (Sigma-Aldrich, St Louis, MO, USA). HRP activity was determined by incubation with TMB (Sigma-Aldrich, St Louis, MO, USA) and H_2_O_2_ at room temperature for 1–10 min and quenching with 1 M H_3_PO_4_. The color was detected using a Biotek Synergy Mx plate reader (Agilent, Santa Clara, CA, USA) at 450 nm.

### Relative and absolute quantities of *N*-glycans

The relative quantity (%) of each *N*-glycan was calculated from the sum of the individual UPLC peak areas in the chromatogram^100^. Each chromatogram area was generated using LC-ESI-HCD-MS/MS, and the quantity (%) of each *N*-glycan (>0.1 %) was determined relative to the total amount of *N*-glycans (100 %)^100^. The absolute quantity of non-overlapping glycan peaks was estimated from the fluorescence intensity of the glycan peak in the UPLC chromatogram, using a linear calibration curve (r2 = 0.99) generated from various concentrations of AB or ProA. The quantities of the remaining glycans were calculated using the relative quantities of fetuin and IgG from the absolute quantity represented by the glycan peak^100^.

### Analysis of UPLC chromatograms

The main glycan peaks were identified and annotated for each profile. The total abundance fraction of each *N*-glycan was calculated from the fraction of individual UPLC peak areas relative to the total area. Owing to some unannotated peaks, most samples did not reach a summed abundance of 1. Some poorly annotated samples identified as outliers by the Matplotlib boxplot^101^ were excluded. We normalized the rest of the glycoprofile dataset for each sample to obtain a total abundance of 1. The raw UPLC peak images provided only the overall glycan composition. Therefore, detailed structures and glycosidic linkages were derived based on references from the literature^74,102^. Finally, we converted the glycan annotation into linear code^52^ style for our choice of nomenclature.

### Analysis of ELLA binding profiles

The assay signal for each lectin was calculated as the average background-subtracted intensity for each sample. Subsequently, the data for each sample were normalized to a total abundance of 1.

### Lectin profile simulation

Based on the literature, we selected eight lectins that mutually recognize ten distinctive common *N*-linked glycan features (Table S1). These eight lectins were selected based on two critical factors. First, the selected set of lectins should be able to recognize most of the *N*-linked glycans found in the glycoprofile dataset. Second, these lectins exhibit high specificity and affinity towards their intended glycan epitopes.

For any given glycoprofile (GP), lectin profile (LP) can be generated by using Equations 1 and 2 below.

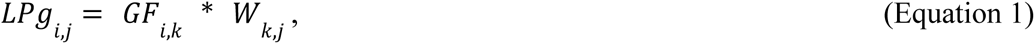

*LPg*_*i,j*_ is the lectin binding profiles for given glycans, where each row represents a glycan and each column represents a lectin; *GF*_*i,k*_ indicates the expressed glycan feature *k* on glycan *i*; and *W*_*k,j*_ is lectin binding rules that tells the frequency of glycan feature *k* recognized by lectin *j*. In this study, we assume that glycan features are recognized by their occurrence counts. Binding rules were adapted from literature^48^, where they provided p-values for significant epitopes bound by each lectin. We converted the p values to z-score, and used it as our binding rule matrix (see Table S3).

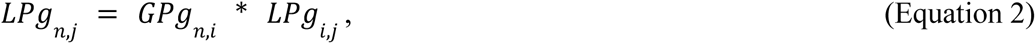

*LPg*_*n,j*_ is the lectin binding profiles for given glycoprofiles, where each row represents a specific glycoprofile, and each column represents a lectin; *GPg*_*n,i*_ is the signal intensity (relative MS/HPLC/UPLC relative area under peak) of glycan *i* in the given glycoprofile *k*.

To calibrate the simulation, we performed regression fitting using the sklearn.linear_model.LinearRegression function. Regression analysis was used to calculate the estimated coefficients and intercepts for each lectin using the measured and simulated lectin profiles from our IgG and Fetuin B samples. Finally, we applied this linear fitting to *LPg*_*n,j*_ to correct the lectin profile simulation..

To demonstrate the problem of mapping a single lectin binding profile to a specific glycoprofile, we generated random glycoprofiles and simulated lectin profiles until we obtained 100 highly correlated profiles (with a Pearson R-correlation above 0.95) from the same glycoprofile. This requires the generation of 2912 different glycoprofiles.

### Model tuning and training

Using lectin profile data as input, where lectins were represented as features on the rows, we constructed a neural network model that produced multiple continuous values. Each value corresponds to the relative abundance fraction of glycans. Model training was performed using the TensorFlow framework. We used RMSprop as the optimizer and softmax activation as the output layer. We trained the model with 60% of our training data at a batch size of four for 200 epochs using the KerasRegressor and GridSearchCV functions to fine-tune the hyperparameters. This grid search method tested every possible combination within predefined hyperparameter ranges (Table S2) to identify the set that yielded the best model performance. To ensure a robust evaluation and mitigate the risk of overfitting, we also utilized 5-fold cross-validation during the tuning phase. The dataset was partitioned into five distinct subsets, with the model trained on four of these subsets and validated on the remaining subset in a cyclic manner. This procedure was repeated five times to ensure that each subset served as the validation set. The performance metric negative RMSE, averaged over five folds, was then used to determine the optimal hyperparameter combination. This process was repeated ten times to evaluate the average performance. The model consisted of four hidden layers, and 20 nodes performed best (Table S2).

When training the model, we used the full set of training data to test IgG and Fetuin B samples. We used MSE as the loss function and executed 1000 epochs. For each experiment, training and testing were repeated 10 times. The average output was calculated and reported.

### Dirichlet distribution to simulate Fetuin B glycoprofiles

The Dirichlet distribution (DD) was motivated by the alignment between the statistical properties of the DD and the normalization requirements (i.e., all glycan abundances sum up to 1) of our glycoprofile data. This allows flexible modulation of glycan abundance via parameterization of the distribution, allowing us to determine their dispersion or uniform distribution in accordance with our specific objectives. To simulate Fetuin B glycoprofiles, we selected a set of eight glycans encompassing various isomeric forms that were previously identified from our Fetuin B glycoprofiling data. Because the abundance of each glycan was unknown, we evenly set the parameters for the distribution at the beginning. Thus, an array of eight equal elements was input into the Numpy.random.dirichlet function, and 30 results were output with different random seeds, as illustrated in Fig. 5A. The generated 30 synthetic glycoprofiles were used for further analysis.

### SHapley Additive explanations analysis

We performed SHapley Additive explanations (SHAP) analysis using the KernelExplainer function from SHAP library (version 0.41.0)^47^. SHAP provides local interpretation, so we randomly selected one sample from the IgG and Fetuin B data for analysis. SHAP values of all the eight lectins were determined. As we trained and tested ten replicate models, the SHAP values are averages derived from all ten models. These SHAP values, either positive or negative, delineate the extent of their impact within the model. This relationship between the relative abundance of certain lectin-glycan interactions in a given lectin profile and their correlation with a given *N*-glycan structure is effectively illustrated by the corresponding bipartite network diagrams (Fig. 3C and 4C).

## Abbreviations

CHO: Chinese Hamster Ovary
geCHO: glyco-engineered Chinese Hamster Ovary
simLP: simulated Lectin Profile
expLP: experimental Lectin Profile
IgG: Immunoglobulin G
DD: Dirichlet Distribution
SHAP: SHapley Additive exPlanations
ex-GPs: experimentally-observed glycoprofiles
comb-GPs: a combined simulated/experimental glycoprofiles

## Author information

**H.L., A.G.P., A.W.T.C.,** and **N.E.L.** conceived the work, analyzed the data and drafted the manuscript. **A.G.P.** and **J.S.** designed and performed the lectin binding experiments on IgG and Fetuin B. **H.L.** curated the measured glycoprofile and lectin profile datasets and simulated lectin binding profiles. **H.L.** and **S-M.S.** implemented the model architecture. **B.C.** and **M.M.** performed the *N*-glycan structure and composition analysis of IgG and Fetuin B. **H.L.** trained the model and performed the analyses on the datasets of IgG and Fetuin B. **H.L.** interpreted the model with SHAP analyses. **J.A., A.M.H, S.S.,** and **B.V.** produced recombinant proteins in CHO cells and determined their *N*-glycan structures.

## Supplementary Information

**Figure S1.**
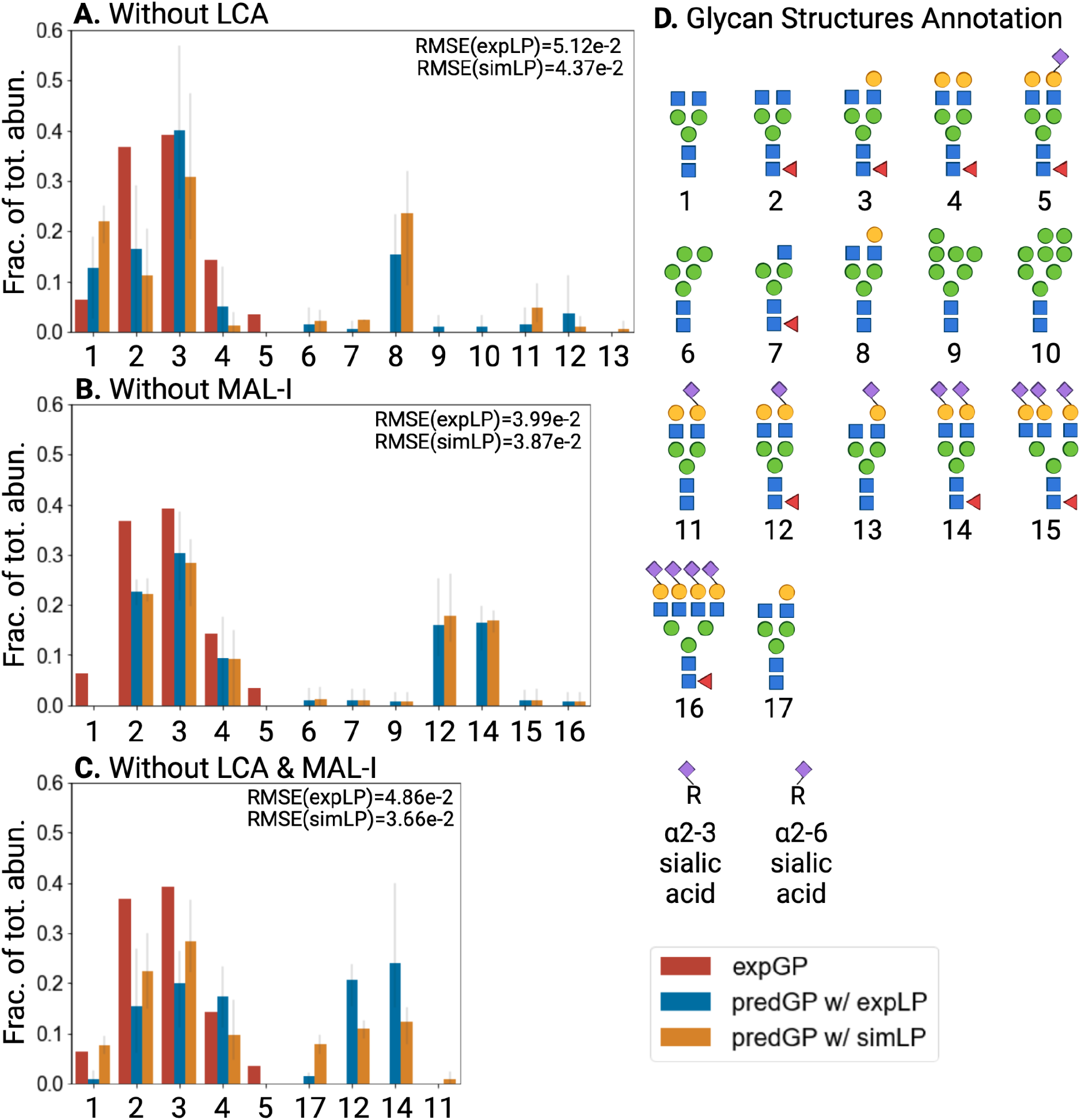
IgG glycoprofile predictions when trained without LCA or/and MAL-I. **A)** Figure S1. IgG glycoprofile predictions trained without LCA and/or MAL-I. A) When LCA was excluded, the model produced a significant false-positive prediction of glycan 8, which does not express core fucose. **B)** When MAL-I is excluded, the model produces two false positive predictions for glycan 12 and 14, which are sialylated. **C)** When LCA and MAL-I are both excluded, the model produces false positive predictions on glycan 12, 14, and 17, which are either sialylated or core-fucosylated.

**Table S1.**
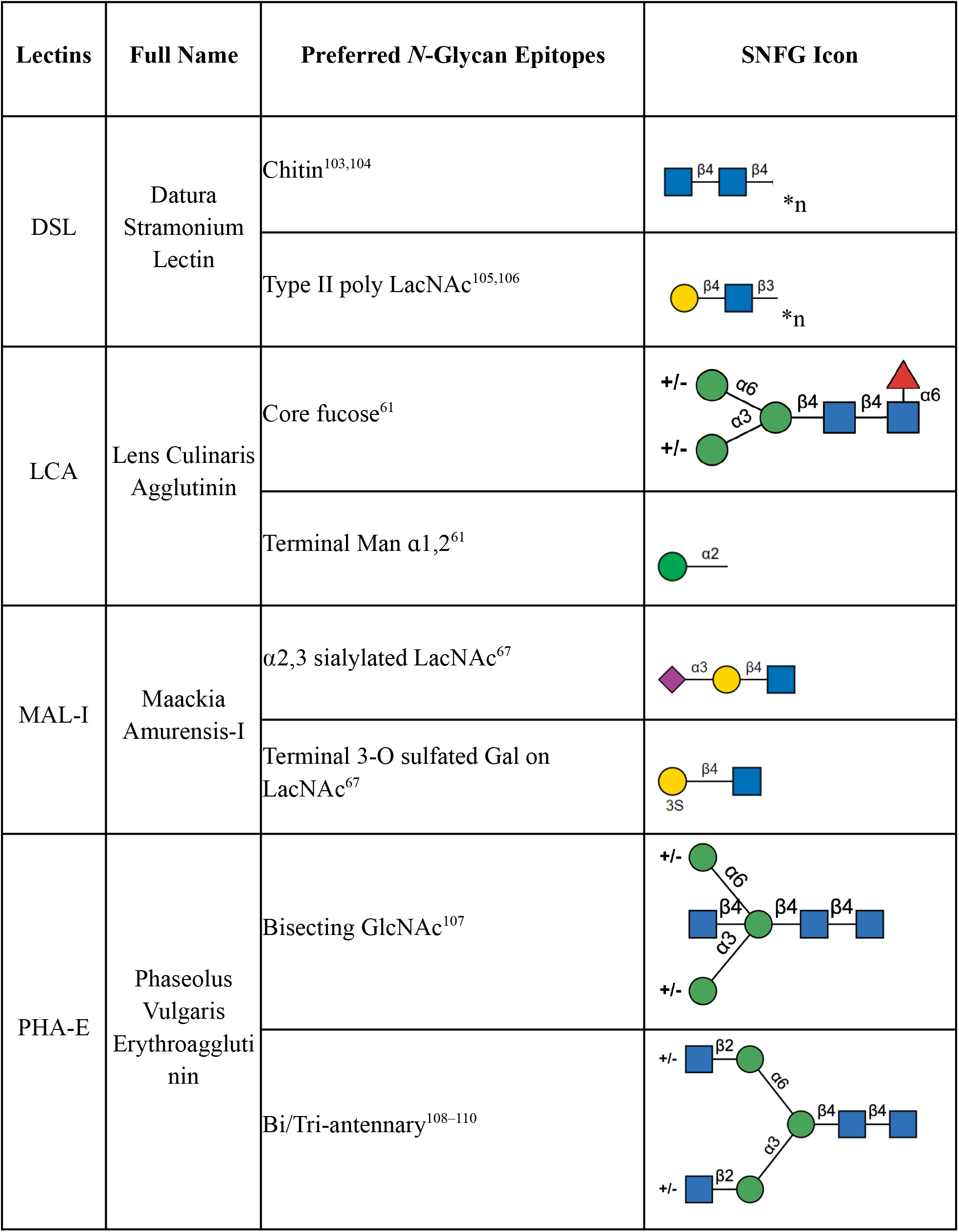
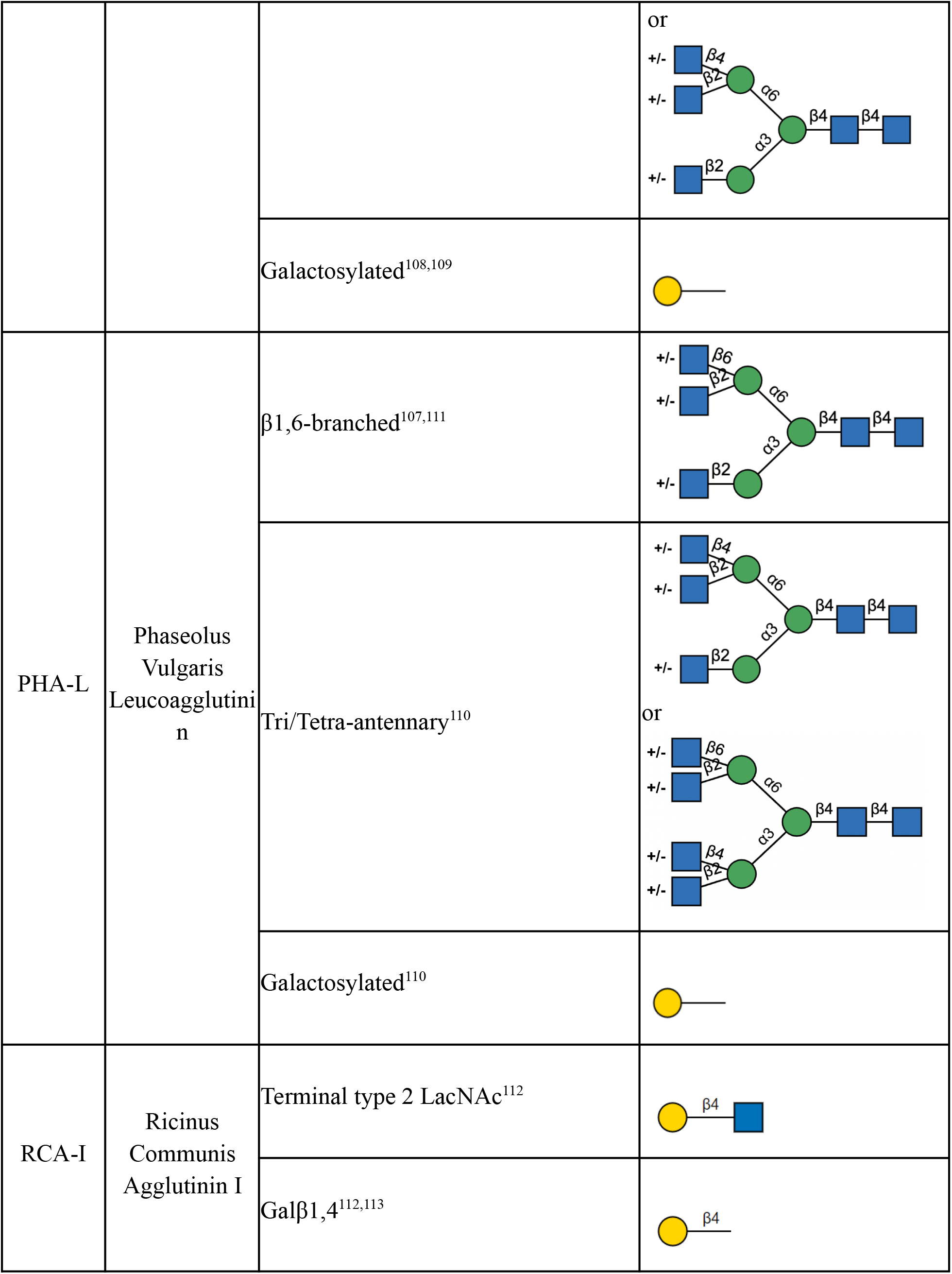
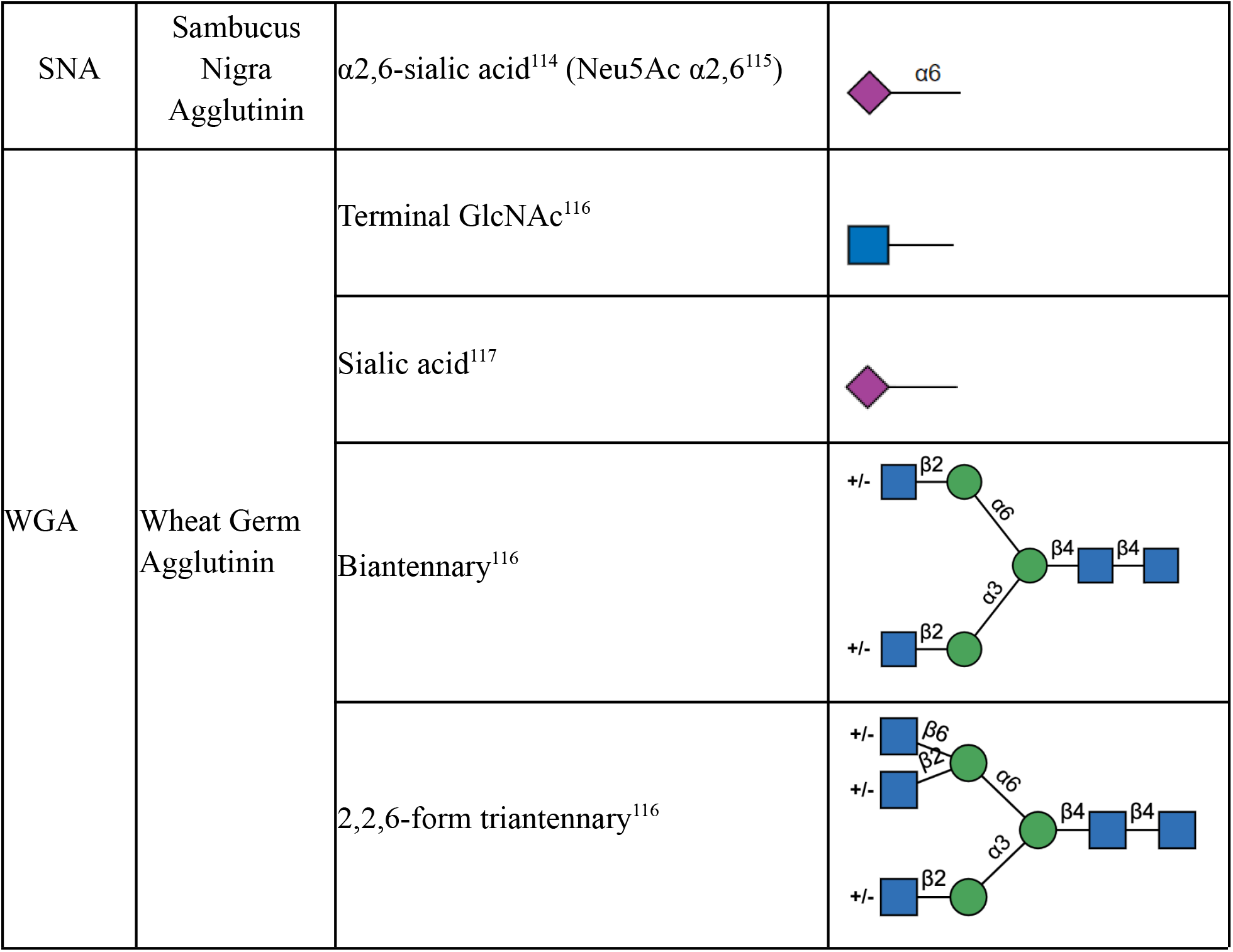
Selected lectins for *N*-glycan lectin profiling.

| Lectins | Full Name | Preferred *N*-Glycan Epitopes | SNFG Icon |
| --- | --- | --- | --- |
| DSL | Datura Stramonium Lectin | Chitin<sup>103,104</sup> |  |
|  |  | Type II poly LacNAc<sup>105,106</sup> |  |
| LCA | Lens Culinaris Agglutinin | Core fucose<sup>61</sup> |  |
|  |  | Terminal Man alpha1,2<sup>61</sup> |  |
| MAL-I | Maackia Amurensis-I | alpha2,3 sialylated LacNAc<sup>67</sup> |  |
|  |  | Terminal 3-O sulfated Gal on LacNAc<sup>67</sup> |  |
| PHA-E | Phaseolus Vulgaris Erythroagglutinin | Bisecting GlcNAc<sup>107</sup> |  |
|  |  | Bi/Tri-antennary<sup>108-110</sup> |  |
|  |  |  | or |
|  |  | Galactosylated<sup>108,109</sup> |  |
| PHA-L | Phaseolus<br>Vulgaris<br>Leucoagglutinin | β1,6-branched<sup>107,111</sup> | |
|  |  | Tri/Tetra-antennary<sup>110</sup> | or |
|  |  | Galactosylated<sup>110</sup> |  |
| RCA-I | Ricinus<br>Communis<br>Agglutinin I | Terminal type 2 LacNAc<sup>112</sup> |  |
| | | Galβ1,4<sup>112,113</sup> | |
| SNA | Sambucus Nigra Agglutinin | α2,6-sialic acid<sup>114</sup> (Neu5Ac α2,6<sup>115</sup>) | |
| WGA | Wheat Germ Agglutinin | Terminal GlcNAc<sup>116</sup> |  |
|  |  | Sialic acid<sup>117</sup> |  |
|  |  | Biantennary<sup>116</sup> |  |
|  |  | 2,2,6-form triantennary<sup>116</sup> |  |

**Table S2.** Negative RMSE values of the model when tuning the hyperparameters.

|  |  | Neurons per Layer |  |  |  |
| --- | --- | --- | --- | --- | --- |
|  |  | 5 | 10 | 15 | 20 |
| Number of Layers | 1 | -3.96E-02 | -3.87E-02 | -3.83E-02 | -3.78E-02 |
|  | 2 | -4.01E-02 | -3.79E-02 | -3.72E-02 | -3.68E-02 |
|  | 3 | -4.01E-02 | -3.79E-02 | -3.72E-02 | -3.63E-02 |
|  | 4 | -4.00E-02 | -3.80E-02 | -3.66E-02 | **-3.60E-02** |

## Acknowledgments

The authors would like to thank Leo Alexander Dworkin and Erik N Bergstrom for feedback. This work was supported by funding from the Novo Nordisk Foundation (NNF20SA0066621) and NIGMS (R35 GM119850). Figures are created with BioRender.com.

